## Supplementary Figures for "The Implications of Alternative Splicing Regulation for Maximum Lifespan"

**Supplementary Fig. 1.** Workflow of alternative splicing identification and quantification. (a) nonhuman mammal species. (b) humans. *For mouse samples, gene annotations from GENCODE (Mus_musculus.GRCm39.107) were used for transcript quantification instead of de novo transcriptome assembly.

**Supplementary Fig. 2.** Sequence regions used for identification of homologous alternative splicing. (a) exon skipping (cassette exons); (b) mutually exclusive exons; (c) alternative 5’ splice sites (alternative donors); (d) alternative 3’ splice sites (alternative acceptors); (e) intron retention; (f) alternative first exons; (g) alternative last exons.

**Supplementary Fig. 3.** Seven types of alternative splicing events. (a) AS events within individual species. (b) Conserved AS events with homologs (against mice) in at least 10 species.

**Supplementary Fig. 4.** Type distribution of MLS-associated alternative splicing events

**Supplementary Fig. 5.** Distribution and Q-Q plots of gene expression levels for genes with splicing events positively (Pos) or negatively (Neg) correlated with maximum lifespan in six tissues.

**Supplementary Fig. 6.** Percentage of MLS-associated alternative splicing (AS) events with significant PSI increases across varying ΔPSI thresholds. Significance is determined using a FDR < 0.05 from Fisher's exact tests on median read counts, with lines representing MLS positively correlated, MLS negatively correlated, and background event groups.

**Supplementary Fig. 7.** Type distribution of age-MLS overlapping alternative splicing events.

**Supplementary Fig. 8.** Heatmap showing hierarchical biclustering of age-associated AS events based on RBP motif enrichment. Each row corresponds to a specific AS event, while columns represent the associated RBPs.

**Supplementary Fig. 9.** Heatmap of pairwise tissue similarity for age-associated splicing events that are identified by traditional regression models fitting alternative splicing individually. The similarity score is based on Jaccard index values.

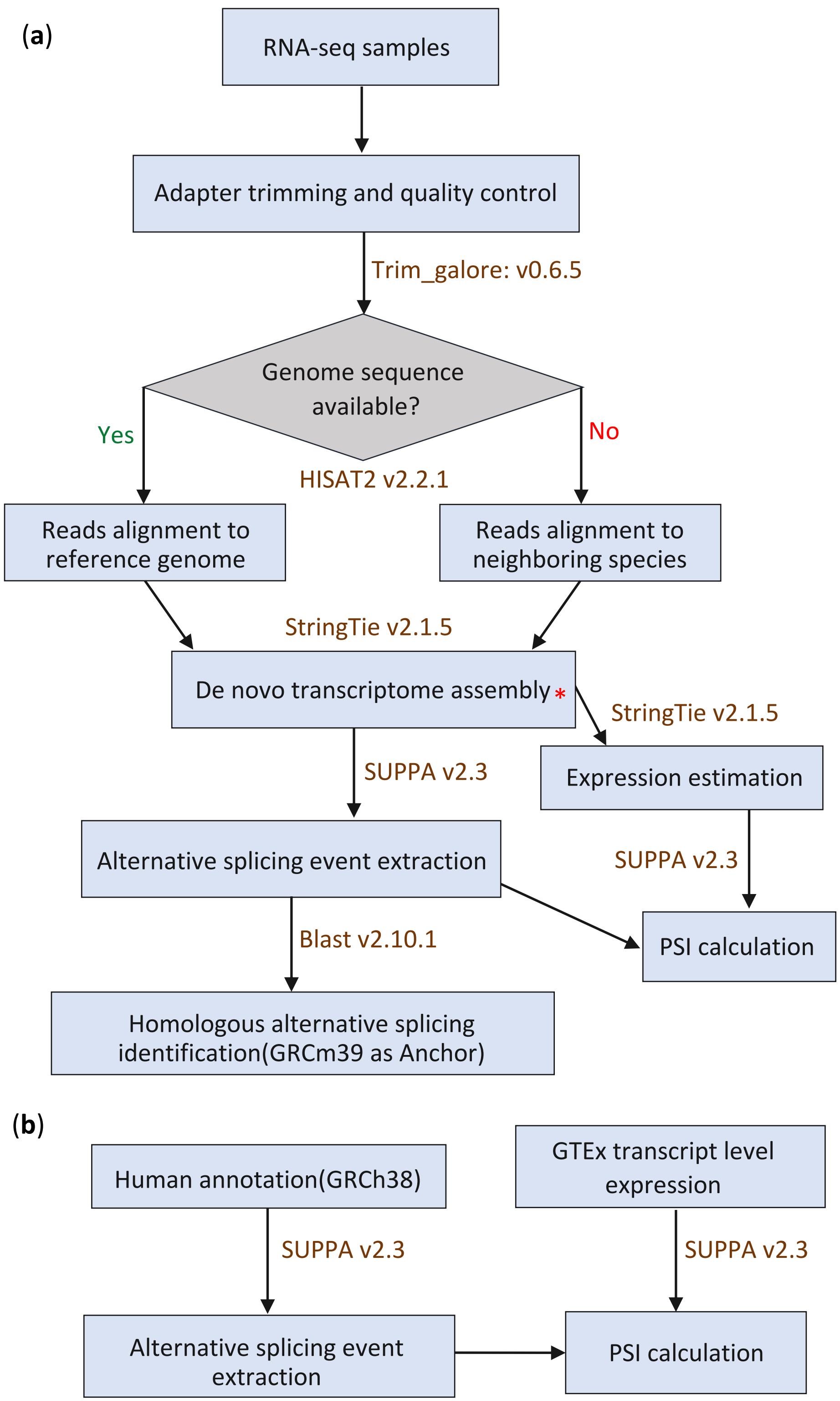


**Supplementary Fig. 1**. **Workflow of alternative splicing identification and quantification.** (a) nonhuman mammal species. **(b)** humans. *For mouse samples, gene annotations from GENCODE (Mus_musculus.GRCm39.107) were used for transcript quantification instead of de novo transcriptome assembly.

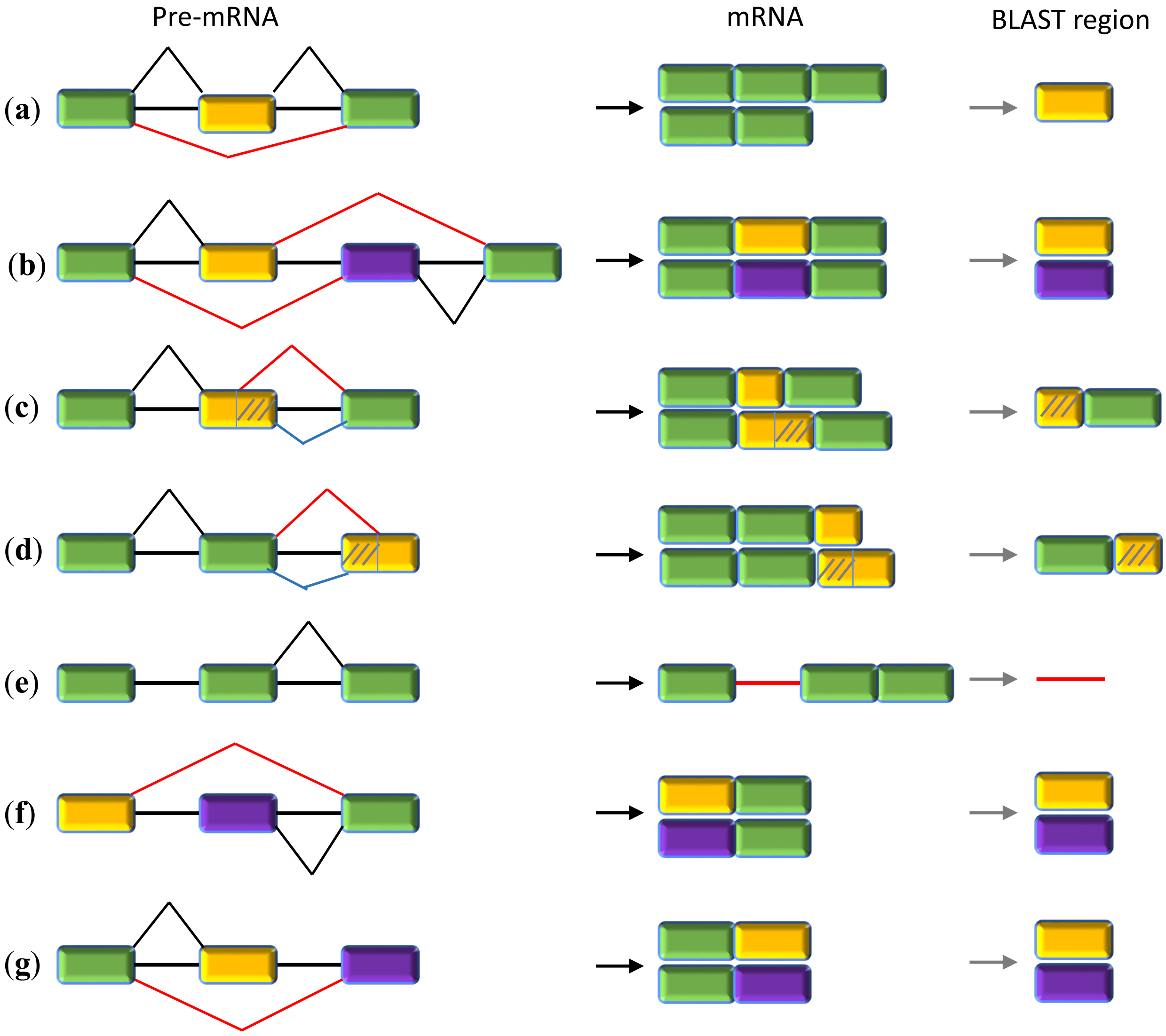


**Supplementary Fig. 2. Sequence regions used for identification of homologous alternative splicing.** (**a**) exon skipping (cassette exons); (**b**) mutually exclusive exons; (**c**) alternative 5' splice sites (alternative donors); (**d**) alternative 3' splice sites (alternative acceptors); (**e**) intron retention; (**f**) alternative first exons; (**g**) alternative last exons.

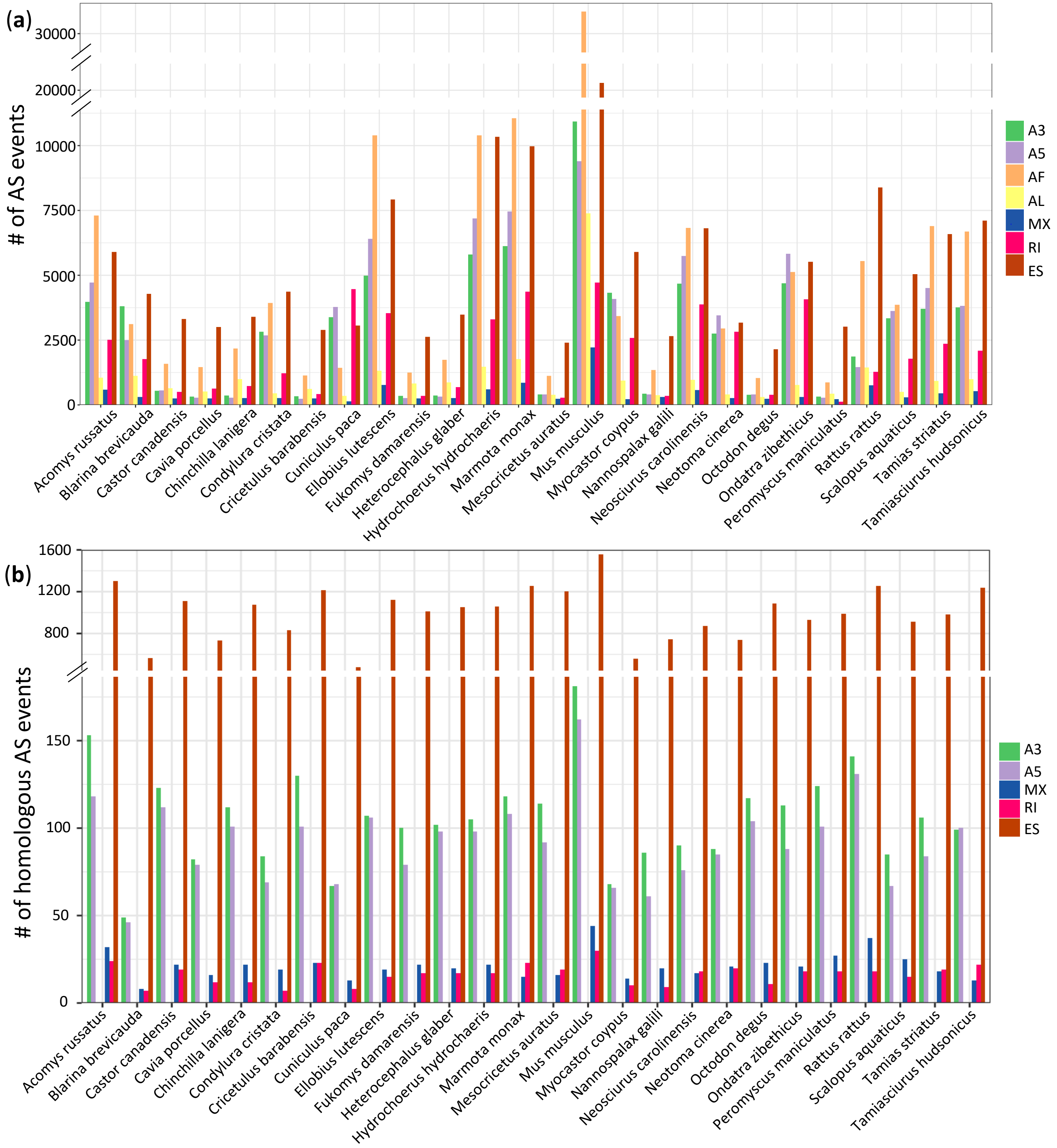


**Supplementary Fig. 3. Seven types of alternative splicing events.** (**a**) AS events within individual species. (**b**) Conserved AS events with homologs (against mice) in at least 10 species.

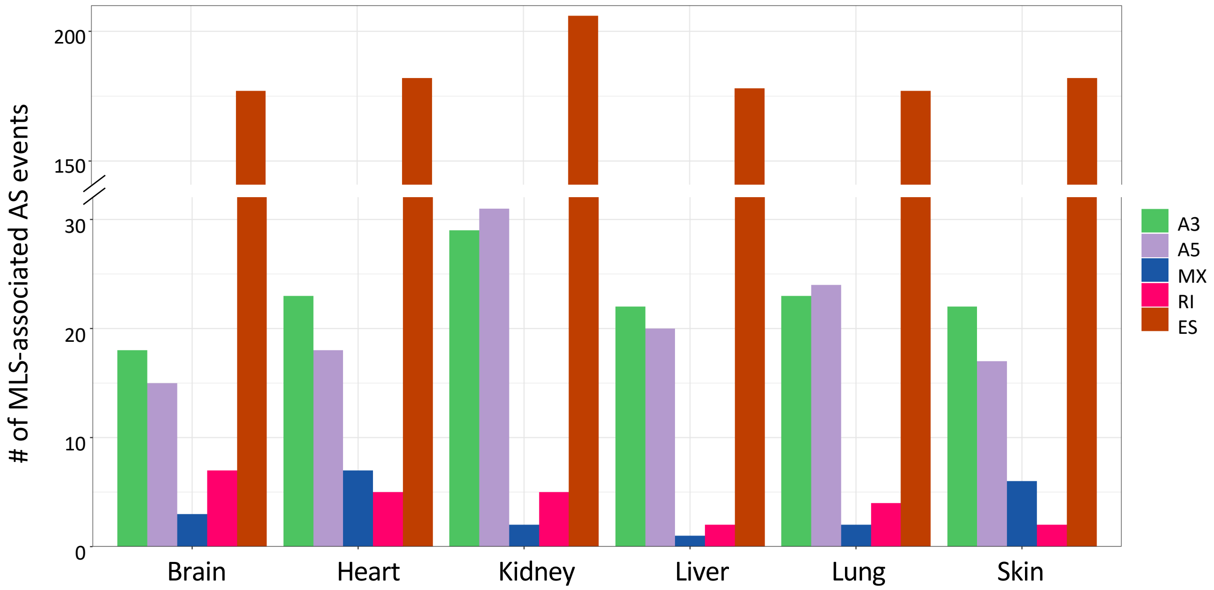


**Supplementary Fig. 4. Type distribution of MLS-associated alternative splicing events.**

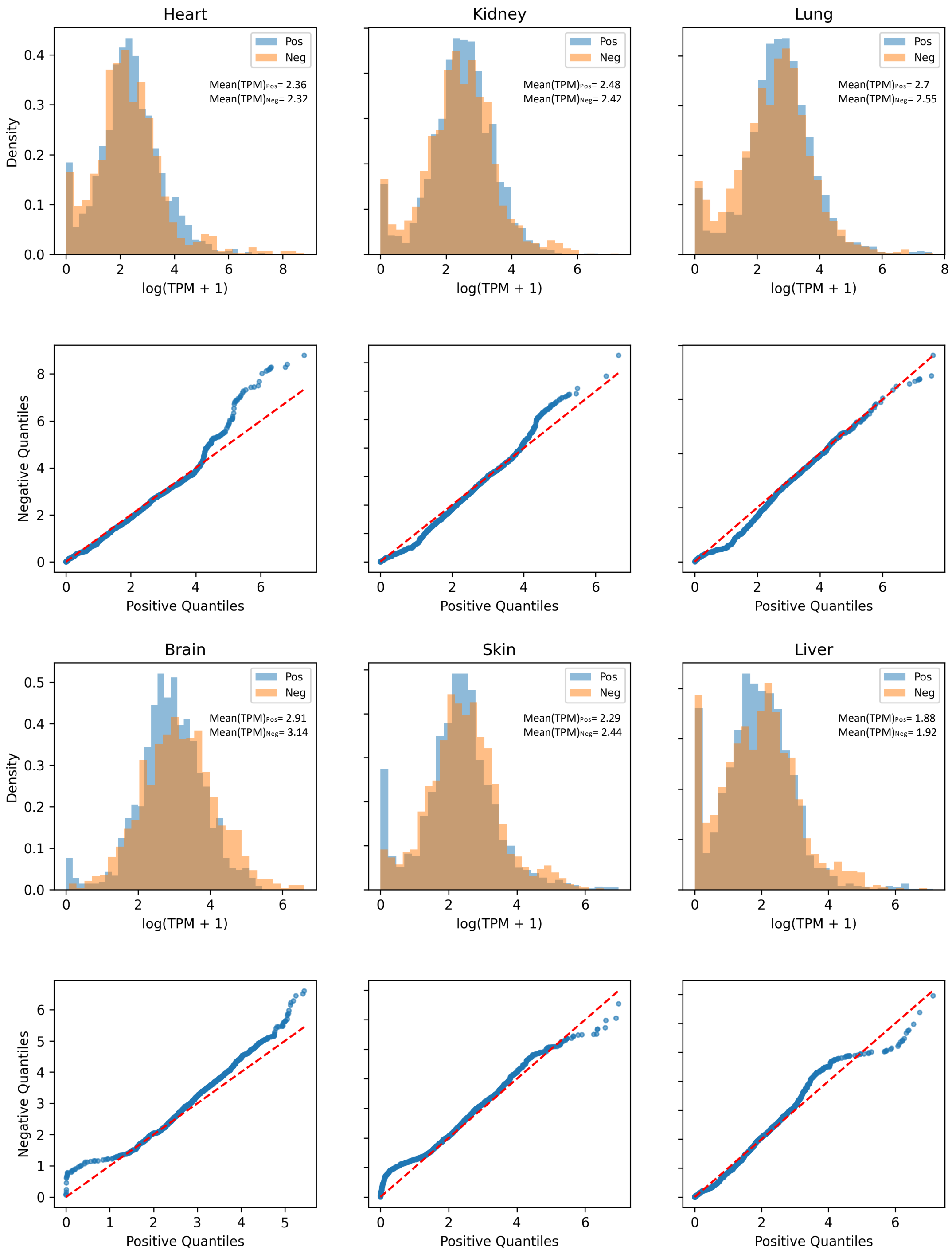


**Supplementary Fig. 5.** Distribution and Q-Q plots of gene expression levels for genes with splicing events positively (Pos) or negatively (Neg) correlated with maximum lifespan in six tissues.

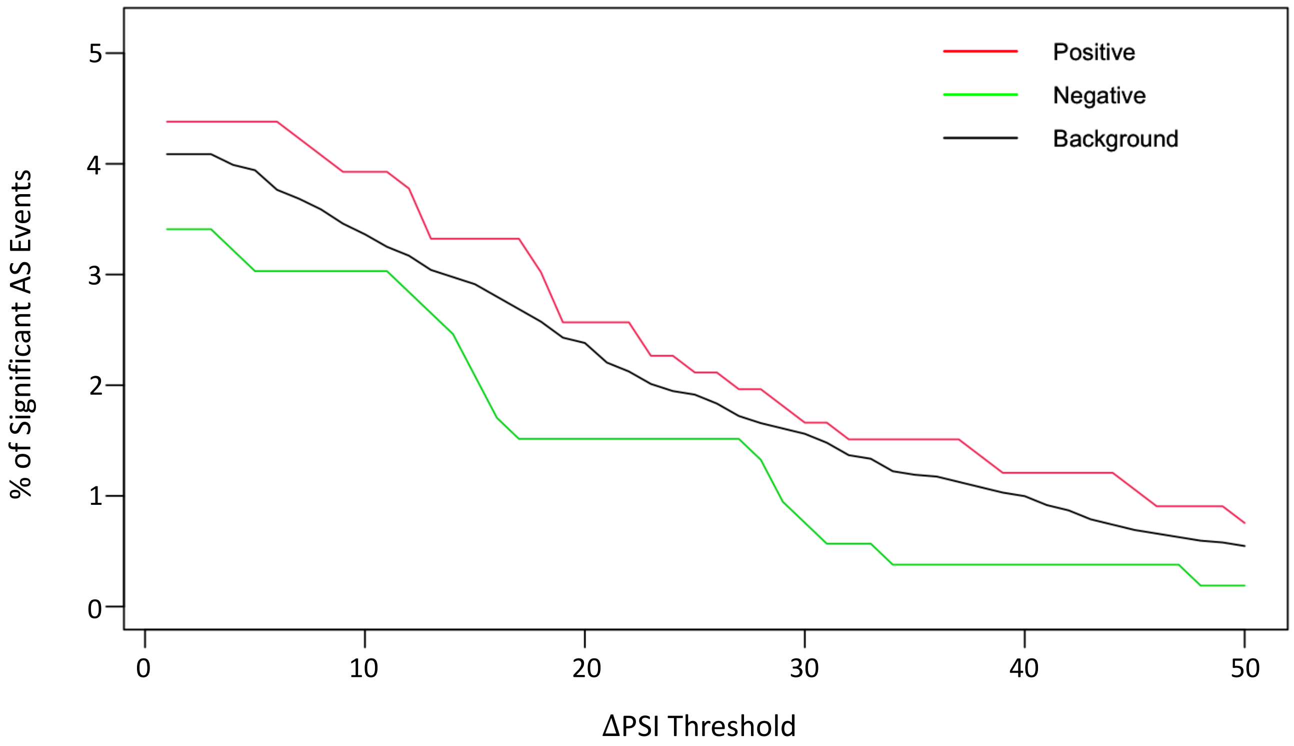


**Supplementary Fig. 6.** Percentage of MLS-associated alternative splicing (AS) events with significant PSI increases across varying ΔPSI thresholds. Significance is determined using a FDR < 0.05 from Fisher's exact tests on median read counts, with lines representing MLS positively correlated, MLS negatively correlated, and background event groups.

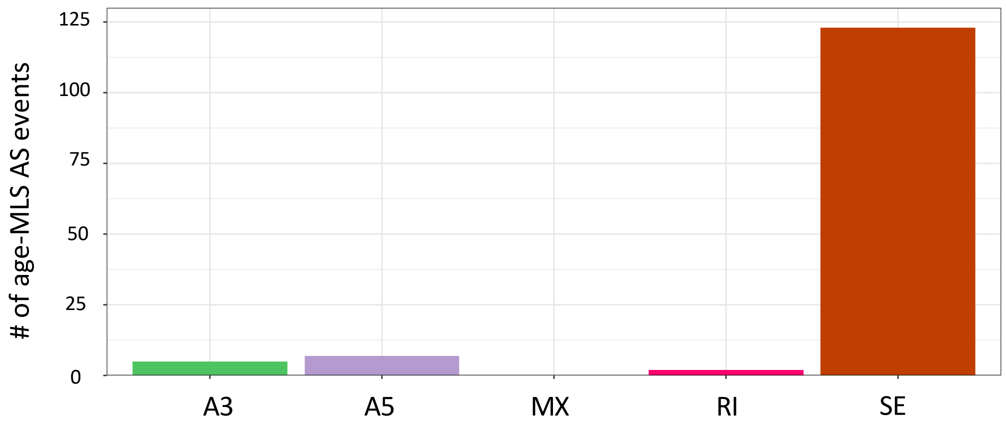


**Supplementary Fig. 7**. **Type distribution of age-MLS overlapping alternative splicing events.**

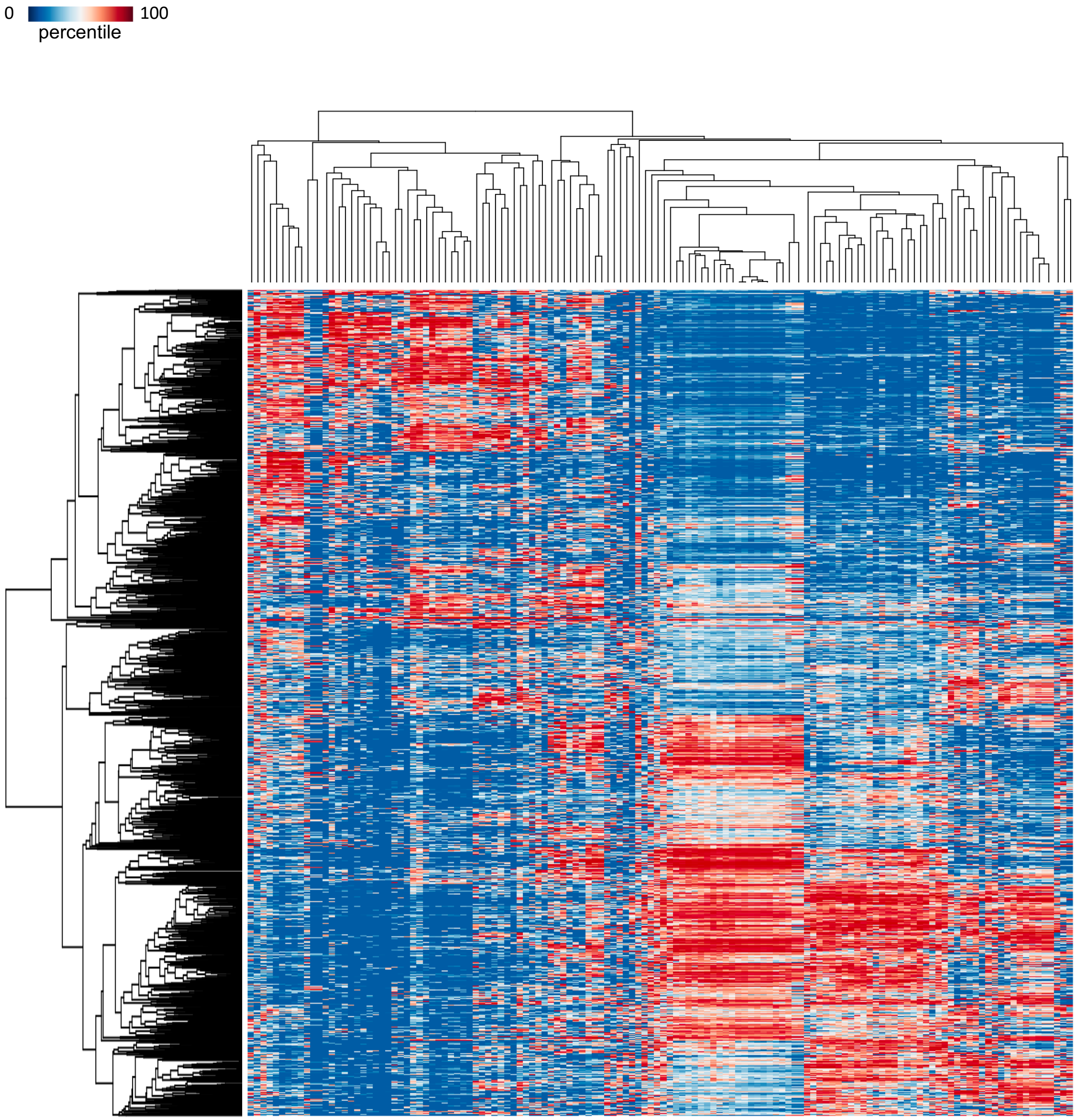


**Supplementary Fig. 8.** Heatmap showing hierarchical biclustering of age-associated AS events based on RBP motif enrichment. Each row corresponds to a specific AS event, while columns represent the associated RBPs.

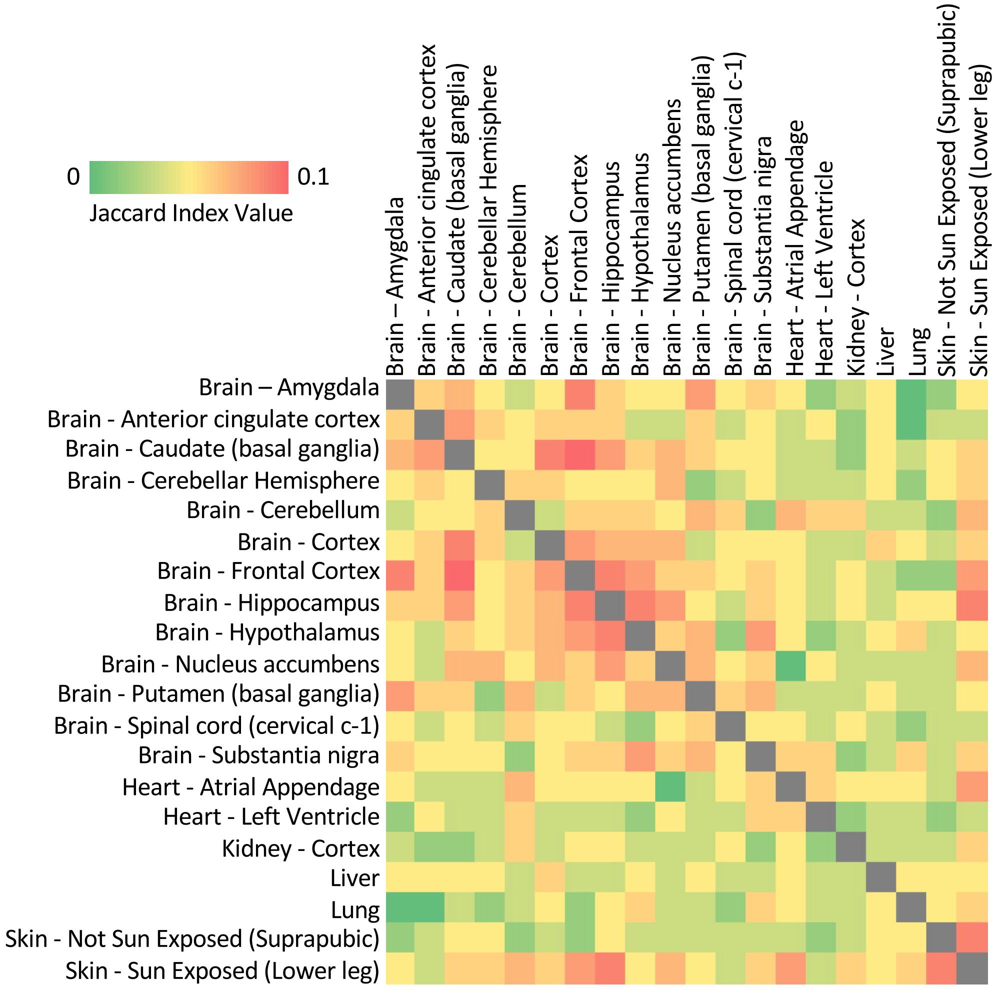


**Supplementary Fig. 9. Heatmap of pairwise tissue similarity for** age-associated **splicing events that are identified by traditional regression models fitting alternative splicing individually.** The similarity score is based on Jaccard index values.
